# Updating contextual sensory expectations for adaptive behaviour

**DOI:** 10.1101/2022.06.08.495309

**Authors:** Ambra Ferrari, David Richter, Floris P. de Lange

**Affiliations:** Radboud University Nijmegen, Donders Institute for Brain, Cognition and Behaviour, 6525 EN Nijmegen, The Netherlands

**Keywords:** Structure learning, statistical learning, sensory expectation, context, belief updating, cortical hierarchy

## Abstract

The brain has the extraordinary capacity to construct predictive models of the environment by internalizing statistical regularities in the sensory inputs. The resulting sensory expectations shape how we perceive and react to the world; at the neural level, this relates to decreased neural responses to expected than unexpected stimuli (‘expectation suppression’). Crucially, expectations may need revision as context changes. However, existing research has often neglected this issue. Further, it is unclear whether contextual revisions apply selectively to expectations relevant to the task at hand, hence serving adaptive behaviour. The present fMRI study examined how contextual visual expectations spread throughout the cortical hierarchy as participants update their beliefs. We created a volatile environment with two state spaces presented over separate contexts and controlled by an independent contextualizing signal. Participants attended a training session before scanning to learn contextual temporal associations among pairs of object images. The fMRI experiment then tested for the emergence of contextual expectation suppression in two separate tasks, respectively with task-relevant and task-irrelevant expectations. Behavioural and neural effects of contextual expectation emerged progressively across the cortical hierarchy as participants attuned themselves to the context: expectation suppression appeared first in the insula, inferior frontal gyrus and posterior parietal cortex, followed by the ventral visual stream, up to early visual cortex. This applied selectively to task-relevant expectations. Taken together, the present results suggest that an insular and frontoparietal executive control network may guide the flexible deployment of contextual sensory expectations for adaptive behaviour in our complex and dynamic world.

**Significance statement:** The world is structured by statistical regularities, which we use to predict the future. This is often accompanied by suppressed neural responses to expected compared with unexpected events (‘expectation suppression’). Crucially, the world is also highly volatile and context-dependent: expected events may become unexpected when the context changes, thus raising the crucial need for belief updating. However, this issue has generally been neglected. By setting up a volatile environment, we show that expectation suppression emerges first in executive control regions, followed by relevant sensory areas, only when observers use their expectations to optimise behaviour. This provides surprising yet clear evidence on how the brain controls the updating of sensory expectations for adaptive behaviour in our ever-changing world.

## Introduction

Extracting regularities from our highly structured environment shapes how we perceive and react to the world (de Lange et al., 2018; Sherman et al., 2020). Sensory expectations let us predict future events and thus react faster and more accurately to expected than unexpected sensory inputs (Kim et al., 2009; Bertels et al., 2012). At the neural level, sensory expectations often generate a decreased neural response to expected than unexpected stimuli, a phenomenon known as ‘expectation suppression’ (Alink et al., 2010; Kaposvari et al., 2018; Richter et al., 2018; Richter and de Lange, 2019; He et al., 2022). Following predictive coding theories (Rao and Ballard, 1999; Friston, 2005, 2010), expectation suppression automatically arises in cortical networks that process the sensory information at hand. For example, predictions about visual object identity produce expectation suppression throughout the ventral visual stream (Richter et al., 2018; Richter and de Lange, 2019; He et al., 2022). Notably, these previous studies show expectation suppression also in a set of downstream regions: insula, inferior frontal gyrus and posterior parietal cortex. However, whether and how their involvement diverges from the one of upstream sensory regions has yet to be addressed, raising questions on the origin of expectation effects in the brain.

Crucially, sensory expectations may need revision as context changes. For instance, whether you are in the Netherlands or in the United Kingdom, you need to pay attention to the same information to drive safely, such as the behaviour of other drivers. However, how events unfold in time depends on the context: entering a roundabout in the Netherlands, you expect vehicles to circulate counter-clockwise and stop on your right, while in the United Kingdom you expect the opposite. As a result, you adapt your behaviour depending on the context. Formally, this process is known as hierarchical structure learning (Collins and Frank, 2013; Gershman, 2017; Collins, 2018): agents organize their state space (e.g. drivers behaviour) into smaller subspaces conditioned on context (e.g. Netherlands or UK) and consequently update their action policy as context changes. While hierarchical structure learning has been extensively investigated during reinforcement learning (Collins, 2018), it has generally been neglected in relation to sensory expectations. Importantly, evaluating their update as context changes may also clarify the role of downstream brain regions. In fact, a frontoparietal executive control network (encompassing inferior frontal gyrus and posterior parietal cortex) is responsible for the selection of task-relevant sensory contingencies in a multidimensional environment, where different contingencies co-exist (Niv et al., 2015; Leong et al., 2017). Similarly, this network may also guide the selection of contextual sensory expectations in a volatile environment, where different contingencies are hierarchically structured by context. Further, such a top-down gating mechanism (Niv et al., 2015; Leong et al., 2017) raises the question of whether contextual updating takes place automatically or rather selectively when expectations are task-relevant, hence serving adaptive behaviour.

The present study examined how sensory expectations spread throughout the cortical hierarchy as participants update their contextual beliefs. We created a volatile environment with two state spaces presented over separate context blocks and controlled by an independent contextualizing signal (Gershman, 2017; Collins, 2018). Participants attended a training session before scanning to learn contextual temporal associations among pairs of object images (Richter et al., 2018). After successful structure learning, corroborated by supplementary behavioural testing, the fMRI experiment tested for the update of contextual expectation suppression under two separate task demands, respectively with task-relevant and task-irrelevant expectations.

In brief, behavioural and neural effects of contextual expectation emerged progressively within a context, accompanied by distinct neural profiles across the cortical hierarchy: expectation suppression appeared first in the insula and frontoparietal regions, followed by the ventral visual stream, up to early visual cortex. This applied selectively to task-relevant expectations. Taken together, these results suggest that an insular and frontoparietal executive control network may guide the flexible deployment of contextual sensory expectations for adaptive behaviour in our complex and ever-changing world.

## Materials and Methods

### Data and code availability

All materials, data and code that are relevant to replicate the current findings are available on the Donders Repository (data.donders.ru.nl) for evaluation during the peer-review process; upon publication, a persistent identifier (DOI) will be made publicly available.

### Participants

The study included the main fMRI experiment and two supplementary behavioural experiments performed via online testing. For the fMRI experiment, a total of 41 volunteers were recruited from the Radboud University research participation system. Following a-priori defined criteria (see section ‘Exclusion criteria - fMRI’), 39 participants underwent MRI scanning and 34 of them were included in the analysis and results (9 males; mean age 26, range 20-40 years). This final number of included participants was based on a-priori power analysis (G*Power 3.1, Faul et al., 2009), computing the required sample size to achieve a power of 0.8 to detect a medium effect size of Cohen’s d = 0.5, at alpha = 0.05, for a two-tailed paired t test.

For the two supplementary behavioural experiments, two independent pools of participants were recruited from Academic Prolific (www.prolific.co). In a structure learning experiment, participants underwent similar tasks to those employed in the main fMRI experiment. In an incidental exposure experiment, we presented the same contextual contingencies unbeknownst to participants, who performed a cover task. In total, we recruited 121 volunteers for the structure learning experiment and 103 volunteers for the incidental exposure experiment. Following a-priori defined criteria (see section ‘Exclusion criteria - Behavioural’), 100 participants were included in the analysis and results of the structure learning experiment (37 males; mean age 29, range 21-40 years) and the incidental exposure experiment (36 males; mean age 30, range 19-40 years). This final number of included participants was based on a-priori power analysis, computing the required sample size to achieve a power of 0.8 to detect a small effect size of Cohen’s d =0.3, at alpha = 0.05, for a two-tailed paired t test. The larger sample size was also motivated by the notion that online studies may produce noisier results than laboratory studies.

All volunteers were right-handed and reported normal or corrected to normal vision and no history of neurological or psychiatric conditions. They provided written informed consent and received financial reimbursement for their participation. The study followed institutional guidelines of the local ethics committee (CMO region Arnhem-Nijmegen, The Netherlands).

### Stimuli

The stimulus material consisted of a pool of 56 full colour images of everyday objects derived from a previous study (Brady et al., 2008). For each participant, we randomly sampled a set of 6 stimuli, thereby eliminating potential effects induced by individual image features at the group level. Objects were either electronic (consisting of electronic components and/or requiring electricity to function) or non-electronic. Stimuli spanned approximately 5°× 5° visual angle and were presented on a mid-gray background in the centre of the screen.

### Experimental design

We used identical designs for the main fMRI experiment and the two supplementary behavioural experiments. In each experimental trial, participants were exposed to two object images presented in quick succession: a leading image was followed by a trailing image (each image duration: 500 ms; no interstimulus interval, ISI). Each stimulus set comprised 6 object images: 2 serving as leading objects and 4 serving as trailing objects (Figure 1A). The expectation manipulation consisted of a repeated pairing of images: each leading image predicted the identity of its two paired trailing images with a 40% chance respectively (expected condition); any of the two other trailing images occurred with 10% chance (unexpected condition). Each trailing image was only (un)expected depending on which leading image preceded it (i.e. P(trailing | leading) = 0.4 for expected condition; P(trailing | leading) =0.1 for unexpected condition). Thus, each trailing image served both as an expected and unexpected image depending on the leading image. In addition, trial order was pseudo-randomized, with different pairs in consecutive trials. In sum, any difference between expected and unexpected trials cannot be explained in terms of individual stimulus frequency or familiarity, adaptation or trial history. Crucially, we created a volatile environment by means of two contexts: the transition probability matrix governing image associations in context 1 was reversed in context 2. The two contexts were presented over separate blocks in alternating order (Figure 1B) and the beginning of each context block was explicitly signalled by a preliminary context cue (see section ‘Experimental procedure - fMRI’). Since we aimed to evaluate how contextual expectation effects emerged as a function of context start, we split each context block into separate consecutive bins.

**Figure 1.**
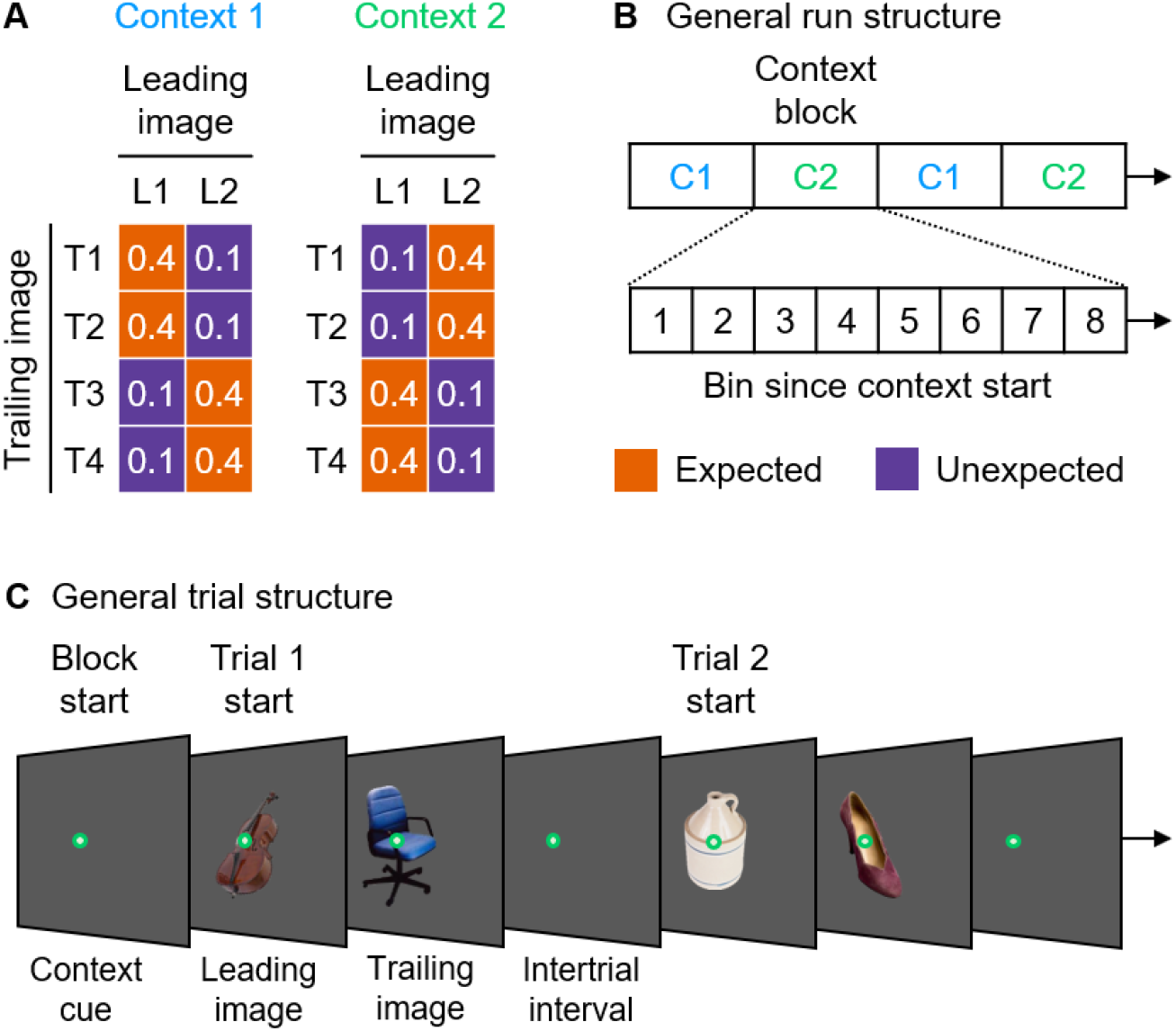
General experimental design and procedure. **A)** Image transition matrix determining image pairs. Two leading images (L1–L2) and four trailing images (T1–T4) were randomly sampled for each participant and session. The expectation manipulation consisted of a repeated pairing of images: each leading image predicted the identity of its two paired trailing images with 40% reliability respectively (expected condition, orange); any other trailing image occurred with 10% reliability (unexpected condition, purple). Further, we created two contexts: the set of image associations presented in context 1 (blue) were reversed in context 2 (green). Thus, the image pairs that were expected in one context became unexpected in the other context and vice versa. **B)** General run structure across tasks. The two contexts (C1 and C2) were presented over blocks of 32 trials (24 expected, 8 unexpected). Each run contained four context blocks (two per type) in alternating order. To evaluate how contextual expectation effects emerged as a function of context start, we created 8 bins for each context block. Each context bin contained 3 expected and 1 unexpected trials. In summary, collapsing across context type, the study conformed to a 2 (expectation: expected / unexpected) × 8 (bin since context start) repeated measures design. **C)** General trial structure across tasks. At the beginning of a context block, a 2-second context cue (i.e., colour of the fixation point) signalled context type. Within a context block, each trial presented a leading image followed by a trailing image (each image duration: 500ms; no ISI). Within the ITI, participants provided their response, depending on the task (see section ‘Experimental procedure - fMRI’).

Each context block was characterized by a 3:1 ratio of expected to unexpected trials (32 trials in total: 24 expected, 8 unexpected). Given this ratio and the constraint of 8 unexpected trials per context block, we created a total of 8 bins per block, with each bin containing 3 expected trials and 1 unexpected trial based on their order of appearance since context start. In other words, we assigned to bin 1 the first 3 trials of the expected condition and the first trial of the unexpected condition, and so forth. This way, we compared expected and unexpected conditions within each bin and we then evaluated how such comparison evolved across bins, since the start of the context. In summary, collapsing across context type, the study conformed to a 2 (expectation: expected / unexpected) × 8 (bin since context start) repeated measures design. Further, to create a general statistical framework across behavioural and fMRI analyses and optimise statistical power for the whole-brain fMRI analysis, we split each context block into two halves (early bins 1:4 / late bins 5:8, see section ‘Statistical analysis - General overview’).

### Experimental procedure - fMRI

The fMRI experiment consisted of two separate sessions per participant on different days. In each session, we sampled a new set of stimuli.

#### Preliminary screening session - day one

The aim of this session was to preselect participants who showed robust structure learning. In a behavioural cubicle, participants performed two tasks. First, they briefly familiarized themselves with the stimuli. In each trial, an object image was presented for 3500 ms in the centre of the screen and participants categorized the object as electronic or non-electronic via two-alternative forced choice (keys counterbalanced across participants). Then, written feedback indicated the correct response (i.e. ‘electronic’ or ‘non-electronic’) for 1500 ms, followed by an intertrial interval (ITI) of 500 ms (6 stimuli × 2 trials / stimulus = 12 trials in total). Second, participants performed a pair recognition task, which evaluated their ability to learn contextual associations among pairs of images. Participants were explicitly instructed to discover which image associations were more likely in each context, while maintaining their gaze on a fixation bull’s-eye (outer circle 0.7° visual angle) that was superimposed on the images in the centre of the screen. At the beginning of each context block (Figure 1C), participants were informed of the context type by a 2-second cue (i.e., colour of the fixation point). Within a context block, each trial presented a leading image followed by a trailing image (each image duration: 500 ms; no interstimulus interval, ISI). Within a 2-second response window since trailing image onset, participants reported whether the trailing image was expected to appear given the leading image via two-alternative forced choice (keys counterbalanced across participants). Then, written feedback revealed the correct response (i.e. ‘expected’ or ‘unexpected’) for 1500 ms, followed by a 500 ms ITI. Each run comprised 4 context blocks (2 blocks per context type presented in alternating order). Participants completed 3 runs, each lasting ~10 minutes ([24 expected trials + 8 unexpected trials] × 2 context types × 2 blocks × 3 runs = 384 trials in total). Contingent on their performance (see section ‘Exclusion criteria - fMRI’), they were invited back for the fMRI session on a different day.

#### fMRI session - day two

Only participants who passed the preliminary screening session participated in the subsequent fMRI session. In a behavioural cubicle outside the scanner, participants performed the same two tasks that they tried during the screening session. First, they briefly familiarized themselves with a new set of stimuli. Second, using this new set of stimuli, they performed the pair recognition task to learn contextual associations among pairs of images. Third, they moved into the MRI scanner, where they were exposed to the same contexts and image pairs. There, they performed two main tasks (i.e. categorization and oddball tasks), in which predictions were respectively task-relevant and task-irrelevant.

In a categorization task, participants reported whether the leading and trailing images belonged to the same category (electronic or non-electronic) via two alternative forced choice (keys counterbalanced across participants). Here, predictions about trailing image identity were task-relevant: knowing which trailing images most likely followed each leading image facilitated the categorization process. In particular, if participants used their predictive knowledge, we expected them to be faster for expected relative to unexpected trailing images (Richter et al., 2018; Richter and de Lange, 2019; He et al., 2022). In line with this prediction, participants were instructed to respond as accurately and fast as possible. Importantly, across the set of 6 images, object category was counterbalanced with respect to image status (leading / trailing), such that participants could not anticipate the correct categorization response solely based on the leading image. Furthermore, object category was also counterbalanced with respect to the expectation condition. Hence, differences in response times (RTs) would not arise by mere response adjustments costs, but instead reflect perceptual surprise to unexpected trailing objects. Please note that, although essential for the categorization task, the above counterbalancing was kept throughout all tasks for consistency. At the beginning of each context block, participants were informed of the context type by a 2-second cue (i.e., colour of the fixation point). Within a context block, each trial presented a leading image followed by a trailing image (each image duration: 500 ms; no ISI). Participants’ categorization responses were collected between the onset of the trailing image and the subsequent ITI (2500– 9500 ms, randomly sampled from a truncated exponential distribution and serving as implicit baseline). Please note that we later limited the behavioural analyses to trials without response outliers (i.e. | RT | > 3 SD above the group mean). Each run comprised 4 context blocks (2 blocks per context type presented in alternating order). Participants completed 2 runs, each lasting ~12 minutes ([24 expected trials + 8 unexpected trials] × 2 context types × 2 blocks × 2 runs = 256 trials in total).

In a separate oddball task, participants pressed a button whenever an image was presented upside down (oddballs: 12.5% of total trials). Oddballs’ appearance was counterbalanced with respect to image status (leading / trailing) and category (electronic / non-electronic). Hence, stimulus predictability was uncorrelated with oddballs’ appearance; in other words, predictions about trailing image identity were task-irrelevant. Otherwise, we used the same trial structure, context blocks, number of trials and runs employed for the categorization task ([24 expected trials + 8 unexpected trials] × 2 context types × 2 blocks × 2 runs = 256 trials in total). Across participants, we counterbalanced the order of categorization and oddball tasks.

After the two main tasks, participants underwent a functional localizer to define individual object-selective brain regions. Participants were exposed to (1) the same object images as shown during the previous tasks and (2) a globally phase-scrambled version of each image (Richter et al., 2018; Richter and de Lange, 2019; He et al., 2022). In a block design, we flashed each image at 2 Hz (300 ms on, 200 ms off) for a period of 11s. To monitor participants’ vigilance, we asked them to press a button whenever an image (object or phase-scrambled object) dimmed in brightness. We inserted 1 baseline block every 6 image blocks. To increase design efficiency, baseline blocks had variable durations (5.5, 8 or 10.5 seconds) presented in counterbalanced order. Participants completed 1 run lasting ~14 minutes (6 images × 2 conditions × 6 blocks = 72 image blocks + 12 baseline blocks = 84 blocks in total).

In all tasks, trial order was pseudo-randomized (localizer: no consecutive trials with same images; all other tasks: no consecutive trials with same image pairs) and response mapping was counterbalanced across participants. Before each task, participants familiarized themselves with the procedure via one preliminary practice run. After scanning, we evaluated whether participants retained the knowledge acquired during the pre-scanning pair recognition task. To this end, we ran a post-scanning pair recognition task that employed the same design and procedure as during the pre-scanning task, but without corrective feedback and with equal conditional probabilities between image pairs to avoid learning at this final test stage. Participants completed 1 run lasting ~3 minutes ([16 expected trials + 16 unexpected trials] × 2 context types = 64 trials in total).

### Experimental procedure - Behavioural

The two supplementary behavioural experiments were performed by two independent pools of participants. Across the two experiments, we used the same tasks that were employed in the fMRI experiment, with the same trial structure (except a fixed ITI = 1500 ms), same context blocks, number of trials and runs. Before starting each task, participants familiarised themselves with the stimuli and procedure via one short practice run. In each experiment, trial order was pseudo-randomized (no consecutive trials with same image pairs) and response mapping was counterbalanced across participants.

#### Structure learning experiment

The aim of this experiment was to verify robust structure learning at the group level. This was an important premise for the fMRI experiment, where we instead pre-selected participants based on the presence of structure learning. In other words, this experiment verified that learning in our experimental setting holds true at the population level, where there is no pre-selection. First, participants performed the same pair recognition task that was presented pre-scanning in the fMRI experiment; thus, they were explicitly instructed to learn which image associations were more likely in each context. Second, participants performed the categorization task. If they used their predictive knowledge, we expected their categorization response to be faster in trials with expected relative to unexpected trailing images. Finally, we evaluated whether participants retained the knowledge acquired during the initial pair recognition task. To this end, we run the same pair recognition task that was employed post-scanning in the fMRI experiment.

#### Incidental exposure experiment

The aim of this experiment was to investigate the presence of incidental, automatic learning in our experimental setting, akin to statistical learning (Saffran et al., 1996; Turk-Browne et al., 2010; Aslin, 2017; Batterink and Paller, 2019; Batterink et al., 2019). If incidental exposure to the contextual regularities triggered automatic learning, the same may have applied during the scanning period in the fMRI experiment. In this case, we would not know whether the fMRI results reflected, at least partially, automatic learning of the contextual contingencies rather than their selective retrial and usage. First, participants performed the categorization task without being informed about the presence of statistical regularities among image pairs, nor about the two different contexts (signalled by the colour of the fixation point as in the other experiments). As a cover story, participants were told that colour changes of the fixation point served as a vigilance task (i.e. press a dedicated button whenever a colour change occurs). This also ensured that participates still paid attention to the colour changes. Incidental learning was probed via comparison of RTs for expected relative to unexpected trailing images. Second, we evaluated whether participants developed any knowledge of the contextual regularities among leading and trailing images. To this end, we run the same pair recognition task that was employed post-scanning in the fMRI experiment.

### Experimental setup - fMRI

The experiment was presented via Presentation 20.2 (Neurobehavioral Systems Inc., neurobs.com; RRID: SCR_002521) on a Windows 7 machine. In the behavioural cubicle outside the scanner, participants sat in front of the computer monitor at a viewing distance of 50 cm and gave responses via a keypad with their right hand. In the MRI scanner, images were displayed via a BOLDscreen 32 (SKU M0135; 1,920 × 1,080 pixels resolution; 60-Hz frame rate) and they were visible to the participants via a mirror mounted on the MR head coil (visual field of ~33° horizontal × ~19° vertical visual angle at a viewing distance of ~109 cm). Participants gave responses via an MR-compatible keypad (Current Designs HHSC-2×4-C) with their right hand.

### Experimental setup - Behavioural

The two supplementary experiments were presented via Gorilla Experiment Builder (Anwyl-Irvine et al., 2020; gorilla.sc). In order to collect precisely timed responses via keyboard presses, we restricted participation to individuals with a computer, as opposed to phones and tablets. Participants were instructed to sit in front of the computer monitor at a viewing distance of 50 cm and to provide responses with their right hand using specific keyboard keys. Specific a-priori exclusion criteria were employed to control for proper task execution (see section ‘Exclusion criteria - Behavioural’).

### fMRI data acquisition

A 3T Siemens Skyra MR scanner was used to acquire both a T1-weighted MP-RAGE anatomical image (TR / TE = 2300 / 3.03 ms, flip angle = 8°, FOV = 256 mm × 225 mm, voxel size = 1 mm isotropic, 192 sagittal slices acquired in sequential ascending direction) and T2*-weighted echoplanar images (EPI) with blood oxygen level–dependent contrast (multiband factor of 4, TR / TE = 1500 / 33.4 ms, flip angle = 75°, BW = 1850 Hz/Px, A/P phase encoding direction, FOV = 192 × 192 × 114 mm^2^, voxel size = 2 mm isotropic, 68 axial slices acquired in interleaved ascending direction). For each participant, we acquired 425 volumes × 2 runs for the categorization and oddball tasks respectively, and 606 volumes × 1 run for the localizer.

### Statistical analysis - General overview

Using RTs and voxel-wise BOLD responses, we investigated how contextual expectation effects emerged within a context. To create a general statistical framework across behavioural and fMRI analyses and optimise statistical power for the whole-brain analysis, we split each context block into two halves (bins 1:4 / bins 5:8). Complementarily, in the ROI analysis we additionally characterized the emergence of expectation suppression across the 8 individual context bins. In the following sections, we provide a detailed description of the behavioural and fMRI analyses. Across both, after testing for normality assumptions, we performed frequentist parametric analyses and we followed up on non-significant results by running the correspondent Bayesian tests (see section ‘Bayesian analysis’). Effect sizes were calculated in terms of partial-eta-squared 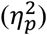 and Cohen’s d_z_ for repeated measures ANOVAs and paired t tests respectively (Lakens, 2013). All standard errors of the mean (SEMs) were calculated as within-subject SEMs. Custom MATLAB 2018b (MathWorks, RRID: SCR_001622) and Python 2.7.13 (Python Software Foundation, RRID: SCR_008394) scripts were used for data handling, statistical tests and data visualization.

### Behavioural analysis

For both the fMRI experiment and the two supplementary behavioural experiments we limited the analyses to trials without missed responses (i.e. no answer within the RT window), premature responses (i.e. RTs < 100 ms) or response outliers (i.e. | RT | > 3 SD above the group mean). Only few trials (across participants’ mean ± SEM) were discarded in the fMRI experiment (4.4% ± 0.1%), in the supplementary structure learning experiment (5.2% ± 0.1%) and in the supplementary incidental exposure experiment (5.5% ± 0.1%). In the following, we describe the behavioural analysis of the three different tasks employed across the fMRI and behavioural experiments.

#### Pair recognition task

Analysis of participants’ responses evaluated whether they possessed robust knowledge of the contextual regularities. We applied the same analysis rationale to the pre- and post-scanning pair recognition tasks using signal detection theory (Wickens, 2002). For each participant, ‘yes’ responses to expected trailing images were labelled as hits, while ‘yes’ responses to unexpected trailing images were labelled as false alarms. It follows that:

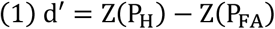

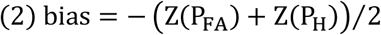

with P_H_ = proportion of hits, P_FA_ = proportion of false alarms. In the preliminary screening session of the fMRI experiment, individual d’ values were used as a criterion to reject participants with subpar performance (see section ‘Exclusion criteria - fMRI’). In the fMRI session and in the supplementary structure learning experiment, individual d’ values were entered into a two-tailed one-sample t test against a predefined threshold (i.e. d’ = 1, consistent with the exclusion criterion) to test for robust knowledge of the contextual regularities at the group level. This represented an important premise for the RT analysis in the following task.

#### Categorization task

Analysis of participants’ RTs evaluated the behavioural effects of contextual expectation on categorisation performance as a function of context start. On each trial, participants reported whether the leading and trailing images belonged to the same category (electronic or non-electronic). If participants relied on learned contextual regularities to provide their categorisation responses, we expected them to be significantly faster in trials with expected relative to unexpected trailing images (Richter et al., 2018; Richter and de Lange, 2019; He et al., 2022). For each participant, median RTs of correct categorization trials were entered into a 2 (expectation: expected / unexpected) × 2 (context half: bins 1:4 / bins 5:8) repeated measures ANOVA. We report two-tailed p-values for the two main effects and the interaction, followed by the planned two-tailed paired t tests evaluating the expectation effect (expected / unexpected) within each context half (bins 1:4 / bins 5:8).

#### Oddball task

Analysis of participants’ responses evaluated vigilance during scanning, in order to reject individuals with subpar performance (see section ‘Exclusion criteria - fMRI’). For each participant, we calculated proportion of hits (i.e. press button after an oddball image, within the ITI) and, complementarily, proportion of omissions (i.e. 1 - proportion of hits).

### fMRI analysis

#### Preprocessing

MRI data were converted to nifti and additional meta data provided, compliant with Brain Imaging Data Structure 1.4.0 (Gorgolewski et al., 2016; bids.neuroimaging.io; RRID: SCR_016124) via BIDScoin 3.6.3 (bidscoin.readthedocs.io). Data quality was subsequently evaluated using MRIQC 0.15.1 (Esteban et al., 2017; mriqc.readthedocs.io). In particular, we quantified instantaneous head motion during scanning in terms of % frame-wise displacement above the critical threshold (default value: 0.2 mm) relative to the full run time series. Data from participants who did not show excessive head motion (see section ‘Exclusion criteria - fMRI’) were included in the remaining analysis steps. Data were preprocessed using fMRIPrep 20.2.3 (Esteban et al., 2019; fmriprep.org; RRID: SCR_016216). Briefly, the fMRIPrep pipeline included brain extraction, motion correction, unwarping and normalization into MNI standard space (only for the whole-brain analysis); finally, FSL FEAT 6.00 (Smith et al., 2004; www.fmrib.ox.ac.uk/fsl; RRID: SCR_002823) was used for spatial smoothing (Gaussian kernel of 5-mm full width at half maximum). The time series of all voxels were high-pass filtered to 1/128 Hz. The first 5 volumes of each functional sequence were discarded to allow for signal stabilization.

#### General linear model (GLM) specification

Voxel-wise GLMs were fit to each participant’s data using FSL FEAT 6.00. The main fMRI tasks (i.e. categorization and oddball) were modelled in an event-related fashion with regressors entered into the design matrix after convolving each unit impulse (duration: 1s, i.e. image pair duration) with a double gamma hemodynamic response function. In addition to modelling the 16 conditions in our 2 (expectation: expected / unexpected) × 8 (bin since context start) repeated measures design, the statistical model included the following nuisance regressors: 6 motion regressors to account for residual motion artefacts; 3 additional nuisance regressors to account for the influence of head motion on the BOLD signal (frame-wise displacement, signal variations in while matter and CSF); context cues (colour change of fixation point at the beginning of each context block); for oddball task only: target trials (upside-down images); first temporal derivative of all regressors. Condition-specific effects (i.e. parameter estimates for the hemodynamic response function regressors) were estimated for each participant according to the general linear model. Data were combined across runs using FSL’s fixed effect analysis.

#### Region of interest (ROI) analysis

Based on previous work (Richter et al., 2018; Richter and de Lange, 2019; He et al., 2022), we preselected early visual cortex (EVC) and object-selective lateral occipital complex (LOC) along the ventral visual stream and superior parietal lobule (SPL), inferior frontal gyrus (IFG) and insula (INS) as downstream regions. Within each ROI (see section ‘ROI definition’), the 16 parameter estimates in our 2 × 8 design were extracted separately from the whole-brain maps. Per participant, the mean parameter estimate of each experimental condition within the ROI was calculated and divided by 100 to yield an approximation of mean percentage signal change relative to baseline. The resulting values were entered into a 2 (expectation: expected / unexpected) × 2 (context half: bins 1:4 / bins 5:8) repeated measures ANOVA, mirroring the RT analysis. We report two-tailed p-values for the two main effects and the interaction, followed by the planned two-tailed paired t tests evaluating the expectation effect (expected / unexpected) within each context half (bins 1:4 / bins 5:8).

Complementarily, we tested for the emergence of expectation suppression over the course of the 8 individual context bins. For each participant and ROI, we first calculated the expectation suppression profile across the context block by taking the unexpected - expected difference of parameter estimates for each of the 8 bins; then, we fitted a linear regression to the resulting expectation suppression profile. A positive regression slope would indicate an increase of expectation suppression along the context block. This was tested for each ROI by subjecting the slope parameters across subjects to a two-tailed one-sample t test, comparing the obtained slopes against zero. Crucially, if downstream executive control regions gated access to the relevant contextual expectations (Niv et al., 2015; Leong et al., 2017), they would show expectation suppression in response to the current set of expected images prior to sensory regions. We thus characterized when we first saw expectation suppression in each ROI by computing the bin value corresponding to the first zero-crossing of the expectation suppression profile (hereafter termed ‘suppression point’). Using a jackknife approach (Miller et al., 1998), a suppression point was computed for each of the n grand average expectation suppression profiles derived from a subsample of n - 1 of the n individual participants (i.e., in a leave-one-out approach, each participant was omitted from one of the subsample grand averages). If an expectation suppression profile (and hence, suppression point) is consistent over participants, then the average value will not change substantially depending on which participant is left out. Please note that this analysis approach does not depend on linearity assumptions for the expectation suppression profiles, thus affording a more precise suppression point estimation. First, for each ROI we tested whether the respective suppression point was significantly different from zero via two-tailed one-sample t tests. Second, to test for pair-wise differences of suppression points between sensory and downstream ROIs, we contrasted the jackknife-estimated suppression points of EVC and LOC respectively with those of SPL, IFG and INS via planned two-tailed paired t tests. Prior to testing for statistical significance, corrected t-values were calculated as follows (Ulrich and Miller, 2001): *t _corrected_* = *t*/(*n* – 1).

#### ROI definition

ROIs were defined bilaterally in each participant’s native space using a combination of anatomical and functional constraints (p < 0.05 at the cluster level Gaussian-random-field (GRF)-corrected within the entire brain, with a cluster-forming uncorrected threshold of p < 0.001). EVC and LOC were defined via the functional localizer, which was modelled in a blocked fashion with regressors entered into the design matrix after convolving each block with a double gamma hemodynamic response function. In addition to modelling the 2 experimental conditions (object / scrambled object), the statistical model included the following nuisance regressors: 9 motion regressors to account for residual motion artefacts (6 motion directions, frame-wise displacement, signal variations in while matter and CSF); target trials (dimmed images). For EVC, the contrast [object + scrambled object] > baseline was constrained to anatomical V1 derived from Freesurfer 6.0.1 cortical surface reconstruction (recon-all; Fischl, 2012; surfer.nmr.mgh.harvard.edu; RRID: SCR_001847). For LOC, the contrast object > scrambled object was constrained to anatomical LOC derived from Freesurfer’s Desikan-Killiany cortical atlas (Desikan et al., 2006). The downstream areas (SPL, IFG, INS) were functionally defined via the group-level contrast ‘unexpected > expected’ derived from a previous study (i.e. categorization task in Richter & de Lange, 2019). The contrast was inverse-normalized to each participant’s native space and constrained to anatomical SPL, IFG and insula derived from Freesurfer’s Desikan-Killiany cortical atlas (Desikan et al., 2006). Finally, all ROIs were constrained to the 300 most active voxels. In order to verify that results did not depend on this arbitrary number of voxels, we repeated all ROI analyses with masks ranging from 100 to 400 voxels in steps of 100 voxels (1562 –6250 mm^3^), in line with previous work (Richter and de Lange, 2019).

#### Whole-brain analysis

We supplemented the ROI analysis with a whole-brain analysis in MNI standard space. For each participant, we evaluated the main effect of expectation (i.e. unexpected > expected and vice versa), the main effect of context half (i.e. bins 5:8 > 1:4 and vice versa) and, crucially, their interaction (i.e. [[unexpected > expected] bins 5:8] > [[unexpected > expected] bins 1:4] and vice versa). In other words, the interaction effect evaluated whether expectation suppression (i.e. unexpected > expected) was stronger in the first or second context half. We further assessed expectation suppression separately in the first half (i.e. [unexpected > expected] bins 1:4) and second half (i.e. [unexpected > expected] bins 5:8). Data were combined across runs using FSL’s fixed effect analysis. FSL’s mixed effect model FLAME 1 was used for the across-participants analysis, which allowed inferences at the population level. We report activations at p < 0.05 at the cluster level GRF-corrected for multiple comparisons within the entire brain, with a cluster-forming uncorrected threshold of p < 0.001.

### Bayesian analysis

To assess whether any non-significant results constituted a likely absence of an effect, or rather indicated a lack of statistical power to detect possible differences, we additionally run the corresponding Bayesian tests. Briefly, all Bayesian analyses were performed using the JZS Bayes factor (Rouder et al., 2009) implemented in JASP 0.11.1 (Love et al., 2019; jasp-stats.org; RRID: SCR_015823) using default settings (Cauchy prior width of 0.707). Interpretations of the resulting Bayes factors quantifying evidence for no effect (B_01_) were based on established classification (Lee and Wagenmakers, 2014).

### Exclusion criteria - fMRI

We excluded participants from all analyses based on three a-priori defined criteria. First, participants were excluded after the preliminary screening session (see section ‘Experimental procedure - fMRI’) if performance during the pair recognition task indicated (1) slack or automatized behaviour, as indexed by excessively fast responses (mean RT < 400 ms) or excessively biased responses (|bias| > 1), or (2) lack of robust knowledge of the contextual regularities (d’ < 1). These exclusion criteria are based on a preliminary pilot study (N = 30). We excluded 2 participants due to lack of robust contextual knowledge. Second, participants were excluded after the fMRI session if performance was subpar during the oddball task (i.e. omissions 3 SD above the group mean) or during the categorization task (proportion of correct responses < 0.85). These exclusion criteria are based on a preliminary pilot study (N = 15). We excluded 1 participant due to subpar performance in the categorization task. Third, we excluded participants who showed excessive head motion during scanning, as indicated by a total % frame-wise displacement (relative motion) 3 SD above the group mean. We excluded 4 participants based on this criterion.

### Exclusion criteria - Behavioural

We excluded participants from all analyses based on two a-priori defined criteria. First, participants were excluded from the structure learning experiment if performance during the pair recognition task indicated slack or automatized behaviour, as indexed by excessively fast responses (mean RT < 400 ms) or excessively biased responses (|bias| > 1). Based on this criterion, 13 participants were excluded from the structure learning experiment. Second, participants were excluded if performance was subpar during the categorization task (proportion of correct responses < 0.85). Based on this criterion, 8 participants were excluded from the structure learning experiment and 3 participants were excluded from the incidental exposure experiment.

## Results

### Successful development and maintenance of contextual expectations

Participants attended a training session before scanning to learn contextual temporal associations among pairs of object images in a pair recognition task. To confirm that they successfully learnt the contextual associations before entering the MRI scanner, we assessed their ability to discriminate expected from unexpected image pairs in each context using signal detection theory. Indeed, d’ in the last run of the pre-scanning pair recognition task was significantly above the threshold value of 1, reflecting robust knowledge of the contextual regularities at the group level (d’ = 3.29 ± 0.13, mean ± SEM; t_(33)_ = 17.77, p < 0.001, d_z_ = 3.05). Likewise, d’ in the post-scanning pair recognition task was significantly above threshold (d’= 2.89 ± 0.14, mean ± SEM; t_(33)_ = 13.48, p < 0.001, d_z_ = 2.31), indicating that participants were able to correctly recall the same contextual associations at the end of the fMRI study, and thus maintained this knowledge during the scanning period. We found consistent results in the supplementary structure learning experiment, where we did not apply any pre-selection criteria for participants inclusion: d’ in the last run of the prescanning pair recognition task was significantly above threshold (d’ = 1.86 ± 0.11, mean ± SEM; t_(99)_ = 7.80, p < 0.001, d_z_ = 0.78). Similarly, participants were also able to correctly retrieve the same contextual associations at the end of the experiment: d’ in the post-scanning pair recognition task was significantly above threshold (d’ = 1.21 ± 0.14, mean ± SEM; t_(99)_ = 8.50, p < 0.001, d_z_ = 0.85). These results confirmed that structure learning in our experimental setting holds true at the population level.

In sum, participants’ performance in the pair recognition task confirmed the successful encoding, storage and retrieval of contextual expectations, which represented an important premise for all subsequent analyses.

### Behavioural effects of contextual expectation develop along the context

Having established robust structure learning, we turned to the behavioural analysis of the categorization task inside the scanner. We anticipated participants to be highly correct at reporting whether the leading and trailing images belonged to the same category (electronic / non-electronic), having pre-selected them based on accurate performance (see section ‘Exclusion criteria - fMRI’). Indeed, the proportion of correct responses was close to ceiling in all experiments (fMRI: 0.95 ± 0.01; supplementary structure learning: 0.95 ± 0.01; supplementary incidental exposure: 0.96 ± 0.01, mean ± SEM). We then concentrated on the RT analysis to assess the behavioural effects of contextual expectation as a function of context start (Figure 2A and Table 1). Participants were significantly faster in trials with expected than unexpected trailing images (i.e. main effect of expectation, F_(1, 33)_ = 14.40, p < 0.001, 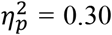). Furthermore, this expectation effect increased over the course of the context block (i.e. expectation × context half interaction, F_(1, 33)_ = 6.77, p = 0.014, 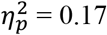).

**Figure 2.**
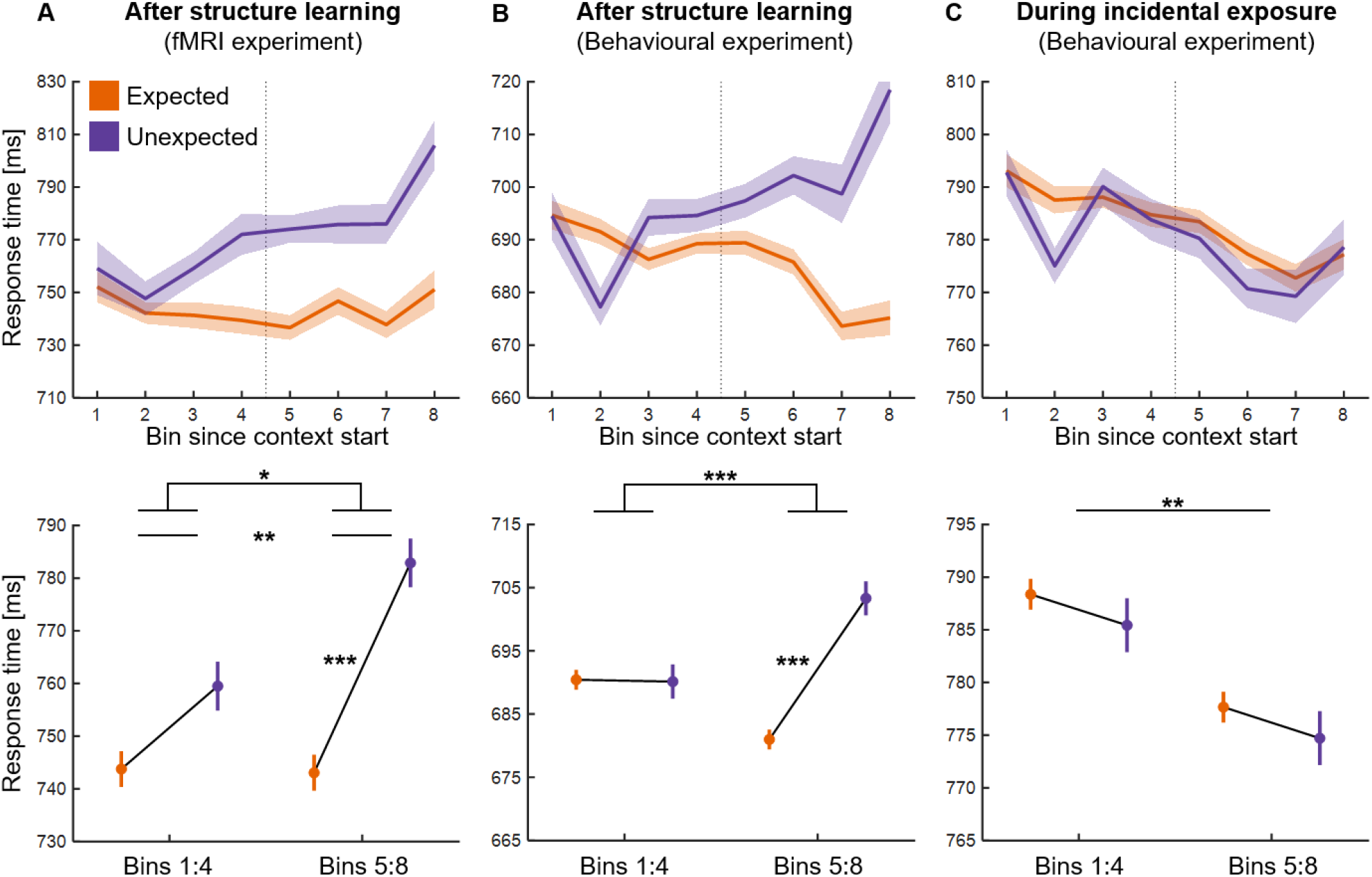
Behavioural effects of contextual expectation in the categorization task. Across participants’ mean (± SEM) response times across the three experiments: **A)** in the fMRI experiment; **B)** in the supplementary structure learning experiment; **C)** in the supplementary incidental exposure experiment. The top row plots data as function of expectation (expected: orange; unexpected: purple) and bin (1-8) since the start of a context block. The bottom row plots the same data as a function of expectation and context half (first half, bins 1:4 / second half, bins 5:8). After structure learning (A-B), participants responded faster in expected relative to unexpected trials in the second context half; such effect was not present during incidental exposure (C). * p < 0.05, ** p < 0.01, *** p < 0.001.

**Table 1.** Results of expectation × context half RT analysis in the categorization task. Main effects and interactions of the 2 (expectation: expected / unexpected) × 2 (context half: bins 1:4 / bins 5:8) repeated measures ANOVA on categorization RTs, after structure learning (fMRI and behavioural experiments) and during incidental exposure (behavioural experiment). Degrees of freedom: d1 = 1, d2 = 33 for the fMRI experiment; d1 = 1, d2 = 99 for the supplementary behavioural experiments.

| Experiment | $F_{(d1,d2)}$ | $p$ | $\eta_p^2$ |
| --- | --- | --- | --- |
| **Structure learning (fMRI)** |  |  |  |
| expectation | 14.40 | < 0.001 | 0.30 |
| context half | 2.87 | 0.100 | 0.08 |
| expectation × context half | 6.77 | 0.014 | 0.17 |
| **Structure learning (Behavioural)** |  |  |  |
| expectation | 14.62 | < 0.001 | 0.13 |
| context half | 0.29 | 0.592 | 0.01 |
| expectation × context half | 16.98 | < 0.001 | 0.15 |
| **Incidental exposure (Behavioural)** |  |  |  |
| expectation | 1.40 | 0.240 | 0.01 |
| context half | 10.30 | 0.002 | 0.10 |
| expectation × context half | 0.00 | 0.996 | 0.00 |

In fact, participants responded significantly faster to expected than unexpected image pairs only in the second context half (t_(33)_ = 4.89, p < 0.001, d_z_ = 0.84), as opposed to the first context half (t_(33)_ = 1.72, p = 0.095, d_z_= 0.29; B_01_ = 1.45).

To corroborate the present findings and address potential alternative interpretations of the results, we turned to the analysis of the two supplementary behavioural experiments. We found consistent RT results in the structure learning experiment, where we did not apply any pre-selection criteria for participants inclusion (Figure 2B and Table 1). A clear expectation effect on RTs was evident only later in the context block (i.e. expectation × context half interaction, F_(1, 99)_ = 16.98, p < 0.001, 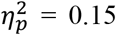). In fact, participants were significantly faster in response to expected than unexpected image pairs only in the second context half (t_(99)_ = 4.95, p < 0.001, d_z_ = 0.49), as opposed to the first context half (t_(99)_ = 0.09, p = 0.931, d_z_ = 0.01; B_01_ = 9.09). In line with the fMRI experiment, this confirms that participants used their predictive knowledge about trailing image identity to speed up their categorization performance, in compliance with task demands. Conversely, we did not observe any expectation effects on RTs in the incidental exposure experiment (Figure 2C and Table 1). Simply, participants became overall faster during the context block (i.e. main effect of context half, F_(1, 99)_ = 10.30, p = 0.002, 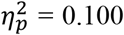). Furthermore, d’ in the subsequent pair recognition task was at chance level, i.e. not significantly different from zero (d’= −0.01 ± 0.02, mean ± SEM; t_(99)_ = −0.19, p = 0.849, d_z_ = −0.02) and with Bayesian results confirming substantial evidence for the absence of an effect (B_01_ = 9.09), suggesting no explicit knowledge of the contextual regularities. Hence, learning was not apparent neither in terms of implicit RT facilitation nor in terms of explicit pair recognition. These results demonstrate that incidental exposure to contextual regularities could not trigger automatic learning, akin to statistical learning (Saffran et al., 1996; Turk-Browne et al., 2010; Aslin, 2017; Batterink and Paller, 2019; Batterink et al., 2019), in the time period afforded by our study, which is in line with the typical duration of statistical learning experiments (Siegelman et al., 2017). In turn, this suggests that expectation effects during the scanning period in the fMRI experiment cannot be explained by automatic learning, but instead reflected the selective retrieval and usage of the pre-established contextual knowledge.

Overall, the RT analysis showed that, after successful structure learning, participants relied on learned contextual regularities to provide their categorisation responses. However, applying this knowledge was not immediate after the start of a context; conversely, behavioural consequences of contextual expectation emerged progressively along the context.

### Neural effects of contextual expectation develop along the context

After successful structure learning, corroborated by supplementary behavioural testing, the fMRI experiment tested for contextual expectation effects at the neural level using voxel-wise BOLD responses. Based on previous work (Richter et al., 2018; Richter and de Lange, 2019; He et al., 2022), we ran an ROI analysis selecting early visual cortex (EVC) and object-selective lateral occipital complex (LOC) along the ventral visual stream and superior parietal lobule (SPL), inferior frontal gyrus (IFG) and insula (INS) as downstream regions. In accordance with the RT results, neural effects of contextual expectation developed over the course of the context block in the categorization task (Figure 3A). In particular, we did not observe any reliable differences between expected and unexpected trials (i.e. main effect of expectation) nor between the first and second halves of the context block (i.e. main effect of context half); instead, we observed a significant interaction between them, reflecting an increase of expectation suppression over the course of the context block. This held true in all the preselected ROIs but EVC, although results appeared qualitatively consistent across all ROIs (Figure 3B and Table 2). Further, the planned paired t tests evaluating the expectation effect within each context half revealed a significantly larger BOLD response to expected than unexpected image pairs in the first context half in LOC (t_(33)_ = 2.56, p = 0.015, d_z_ = 0.44), SPL (t_(33)_ = 2.31, p = 0.027, d_z_ = 0.40) and IFG (t_(33)_ = 2.23, p = 0.032, d_z_ = 0.38), while results remained inconclusive in EVC (t_(33)_ = 1.81, p = 0.079, d_z_ = 0.31, B_01_ = 1.27) and INS (t_(33)_ = 1.35, p = 0.186, d_z_ = 0.23, B_01_ = 2.38).

**Figure 3.**
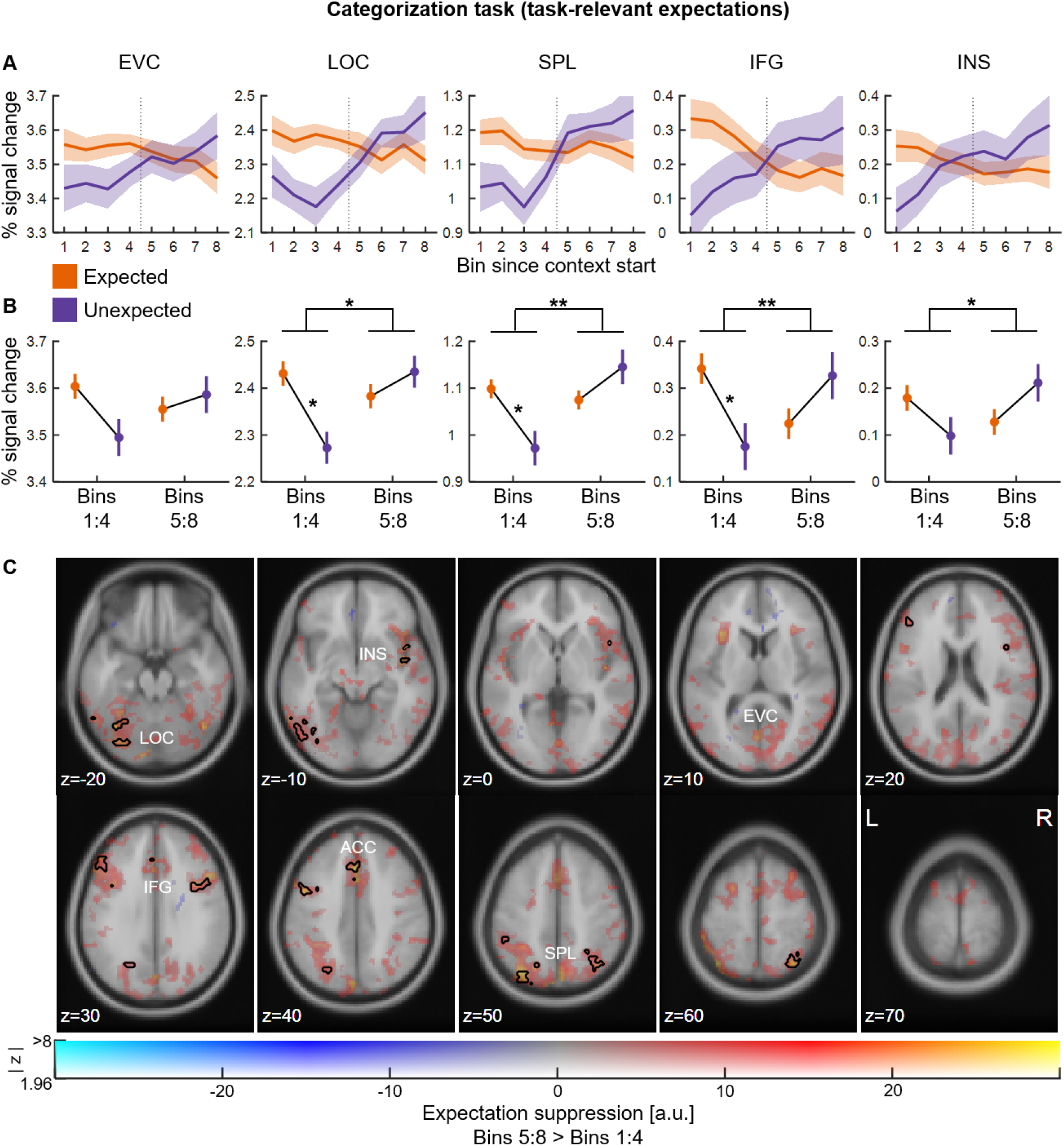
Neural effects of contextual expectation in the categorization task. **A)** Across participants’ mean (± SEM) % signal change relative to baseline for each ROI as function of expectation (expected: orange; unexpected: purple) and bin (1-8) since the start of a context block. **B)** Same data plotted as a function of expectation and context half (first half, bins 1:4 / second half, bins 5:8). **C)** Expectation suppression (i.e. unexpected > expected) in the second context half (bins 5:8) relative to the first context half (bins 1:4), overlaid on the MNI152 2mm template. Colours represent the parameter estimates: red-yellow clusters reflect expectation suppression; opacity represents the associated z statistics. Black contours outline statistically significant clusters (p < 0.05 cluster-corrected). Expectation suppression emerged in the second context half in three widespread networks: ventral visual stream (EVC: early visual cortex, LOC: lateral occipital complex), frontoparietal executive control network (SPL: superior parietal lobule, IFG: inferior frontal gyrus) and salience network (INS: insula, ACC: anterior cingulate cortex). a.u. = arbitrary units, * p < 0.05, ** p < 0.01.

**Table 2.** Results of expectation × context half ROI analysis in the categorization task. Main effects and interactions of the 2 (expectation: expected / unexpected) × 2 (context half: bins 1:4 / bins 5:8) repeated measures ANOVA on % signal change relative to baseline for each ROI: EVC, early visual cortex; LOC, lateral occipital complex; SPL, superior parietal lobule; IFG, inferior frontal gyrus; INS, insula.

| ROI | $F_{(1,33)}$ | $p$ | $\eta_p^2$ |
| --- | --- | --- | --- |
| **EVC** |  |  |  |
| expectation | 1.07 | 0.309 | 0.03 |
| context half | 0.18 | 0.672 | 0.01 |
| expectation × context half | 2.41 | 0.130 | 0.07 |
| **LOC** |  |  |  |
| expectation | 1.48 | 0.233 | 0.043 |
| context half | 1.54 | 0.223 | 0.045 |
| expectation × context half | 7.06 | 0.012 | 0.176 |
| **SPL** |  |  |  |
| expectation | 0.51 | 0.481 | 0.01 |
| context half | 2.43 | 0.129 | 0.07 |
| expectation × context half | 7.80 | 0.009 | 0.19 |
| **IFG** |  |  |  |
| expectation | 0.39 | 0.537 | 0.01 |
| context half | 0.06 | 0.806 | 0.01 |
| expectation × context half | 7.60 | 0.009 | 0.19 |
| **INS** |  |  |  |
| expectation | 0.00 | 0.979 | 0.00 |
| context half | 0.28 | 0.599 | 0.01 |
| expectation × context half | 5.30 | 0.028 | 0.14 |

By contrast, the qualitatively larger BOLD response to unexpected than expected image pairs in the second context half did not reach significance in any ROI (all p > 0.100, B_01_ < 1.80). Here, it is worth noting that expected and unexpected pairs were in fact reversed when the context changed: the pairs that were expected in the previous context became unexpected in the current context. Hence, considering progressive expectation effects in terms of RTs, it is conceivable that what we find in the first context half is in fact a carry-over suppression effect for the expected pairs of the previous block. Over the course of the current block, as participants attuned themselves to the context, BOLD responses to the current expected pairs then became progressively suppressed relative to the current unexpected pairs.

The whole-brain analysis replicated no reliable BOLD response differences between expected and unexpected trials (i.e. main effect of expectation) nor between the first and second halves of the context block (i.e. main effect of context half); instead, we observed an increase of expectation suppression over the course of the context block (i.e. expectation × context half interaction; Figure 3C and Table 3). Cortical areas showing more expectation suppression in the second relative to the first context half included object-selective LOC and fusiform gyrus, superior parietal lobule (SPL) and intraparietal sulcus, inferior frontal gyrus (IFG), insula (INS) and anterior cingulate cortex (ACC; see black contours in Figure 3C). The same contrast also showed the activation of early visual cortex (EVC) at p < 0.001 uncorrected (see red-shaded areas in Figure 3C). Whole-brain results were in line with the ROI analysis also when assessing the expectation effect (i.e. unexpected > expected and vice versa) separately in the first and second context halves. We found a significantly larger BOLD response to expected than unexpected image pairs in the first context half in consistent brain regions to the ones mentioned above, while no significant differences between expected and unexpected trials were present in the second context half (Table 3).

**Table 3.** Results of expectation × context half whole-brain analysis in the categorization task. Listed are significant clusters with their respective area label, MNI coordinates (mm) of the cluster centre of gravity, number of voxels in the cluster, p value of the cluster and its max z statistic. p-values are GRF-corrected at the cluster level for multiple comparisons within the entire brain, with a cluster-forming uncorrected threshold of p < 0.001; exp = expected image pairs; uxp = unexpected image pairs; bins 1:4 = first context half; 5:8 = second context half.

| Contrast | Area label | MNI coordinates |  |  | voxels | p cluster | max z |
| --- | --- | --- | --- | --- | --- | --- | --- |
|  |  | x | y | z |  |  |  |
| [uxp > exp] bins 5:8 ><br>[uxp > exp] bins 1:4 | Superior Parietal Lobule | 33 | -61 | 54 | 234 | < 0.001 | 4.55 |
|  | Intraparietal Sulcus | -28 | -69 | 43 | 216 | < 0.001 | 4.34 |
|  | Lateral Occipital Cortex | -50 | -64 | -13 | 144 | < 0.001 | 4.47 |
|  | Fusiform Gyrus | -31 | -69 | -17 | 163 | < 0.001 | 4.30 |
|  | Inferior Frontal Gyrus | 42 | 8 | 30 | 171 | < 0.001 | 4.39 |
|  | Inferior Frontal Gyrus | -46 | 6 | 36 | 124 | < 0.001 | 4.07 |
|  | Anterior Cingulate Gyrus | -2 | 22 | 36 | 109 | 0.002 | 4.65 |
|  | Supramarginal Gyrus | -48 | 29 | 28 | 161 | < 0.001 | 4.23 |
|  | Insula | 47 | 6 | -8 | 73 | 0.018 | 4.16 |
|  | Inferior Parietal Lobule | -49 | 41 | 49 | 72 | 0.019 | 4.13 |
| [exp > uxp] bins 1:4 | Anterior Cingulate Gyrus | 2 | 9 | 38 | 367 | < 0.001 | 4.47 |
|  | Lateral Occipital Cortex | -54 | -63 | -10 | 157 | < 0.001 | 4.43 |
|  | Fusiform Gyrus | 36 | -77 | -16 | 123 | < 0.001 | 4.43 |
|  | Middle Occipital Gyrus | 33 | -90 | 5 | 121 | < 0.001 | 4.47 |
|  | Middle Occipital Gyrus | -42 | -81 | 15 | 83 | 0.008 | 4.27 |
|  | Supramarginal Gyrus | 64 | -28 | 21 | 278 | < 0.001 | 4.39 |
|  | Supramarginal Gyrus | -61 | -29 | 35 | 78 | 0.011 | 3.98 |
|  | Inferior Frontal Gyrus | 45 | 7 | 30 | 225 | < 0.001 | 4.22 |
|  | Middle Temporal Gyrus | 58 | -60 | 8 | 175 | < 0.001 | 4.25 |
|  | Superior Frontal Gyrus | 28 | 2 | 60 | 199 | < 0.001 | 4.11 |
| [exp > uxp] bins 5:8 | - | - | - | - | - | - | - |
| [uxp > exp] bins 1:4 | - | - | - | - | - | - | - |
| [uxp > exp] bins 5:8 | - | - | - | - | - | - | - |

Combined, the ROI and whole-brain analyses pointed to an increase of expectation suppression for contextually expected pairs within the current context. To characterize this more directly, we complementarily tested for the emergence of expectation suppression (i.e. unexpected > expected) over the course of the 8 individual context bins in all ROIs (Figure 4A). After fitting a linear regression to the expectation suppression profile per participant and ROI, the across-participants’ estimated regression slope ( *b*, mean ± SEM) was significantly above zero for all ROIs: EVC ( *b* = 3.98 ± 1.88, t_(33)_ = 2.12, p = 0.042, d_z_ = 0.36), LOC (b = 4.36 ± 1.65, t_(33)_ = 2.65, p = 0.012, d_z_ = 0.45), SPL (b = 4.15 ± 1.90, t_(33)_ = 2.18, p = 0.036, d_z_ = 0.37), IFG (b = 6.52 ± 2.28, t_(33)_ = 2.86, p = 0.007, d_z_ = 0.49) and INS (b = 4.15 ± 1.65, t_(33)_ = 2.51, p = 0.017, d_z_ = 0.43). These results confirmed an increase of expectation suppression along the context block in all ROIs. To ensure that the present results are not dependent on the arbitrarily selected mask size, we repeated all ROI analyses for mask sizes between 100 and 400 voxels (see section ‘ROI definition’): the direction and statistical significance of all effects were identical across all ROI sizes.

**Figure 4.**
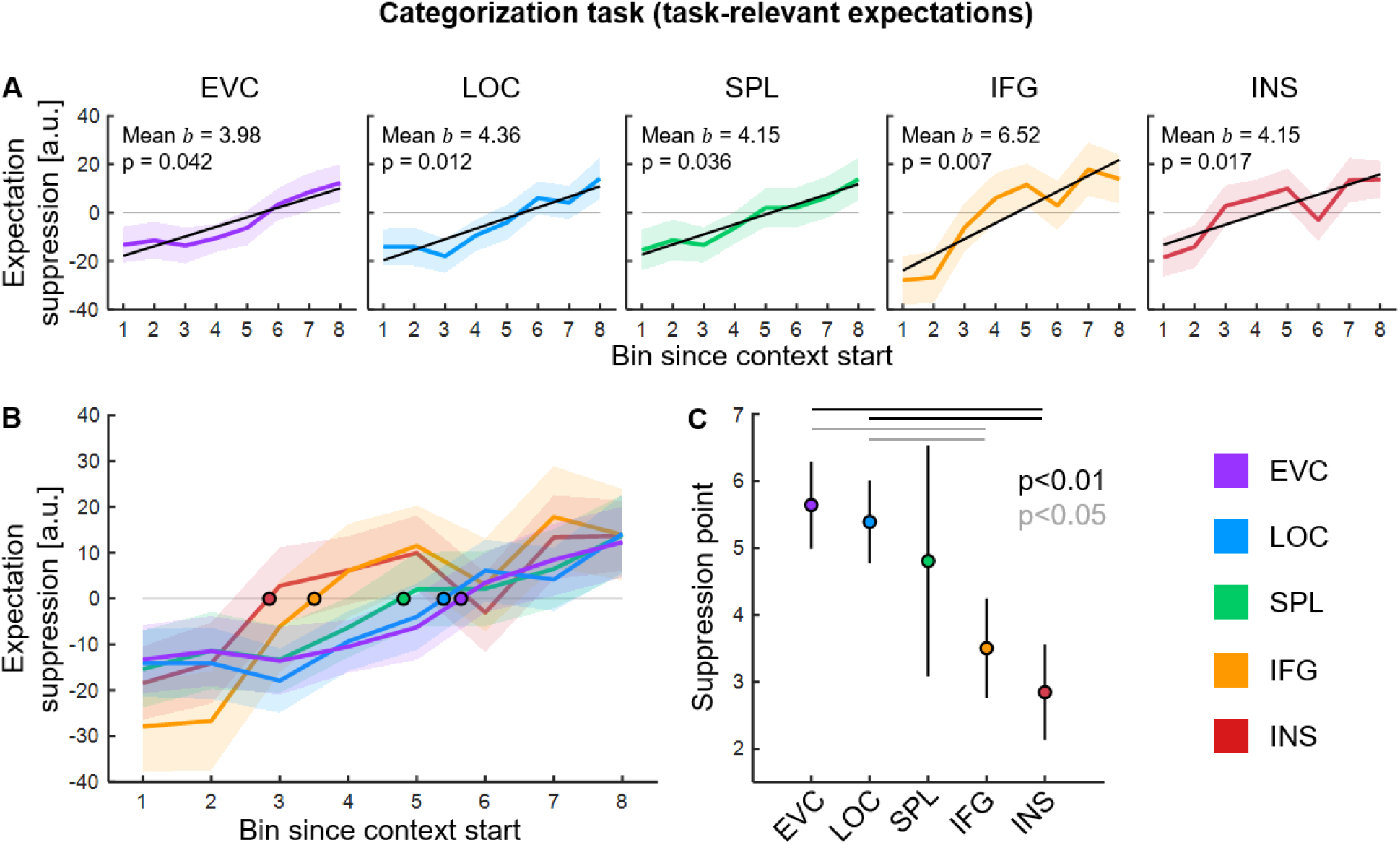
Expectation suppression profile across the cortical hierarchy in the categorization task. **A)** Across participants’ mean (± SEM) expectation suppression profile for each ROI. The regression line in black is derived from the across participants’ mean linear regression parameters (i.e. slope and intercept). The estimated regression slopes ( *b)* were significantly above zero for all ROIs, indicating an increase of expectation suppression over the course of the context block (see across participants’ mean b and p-value in each subplot). **B)** Suppression point for each expectation suppression profile (coloured dot with black contour), showing that expectation suppression emerged at different moments across the cortical hierarchy: first in the insula (INS), followed by inferior frontal gyrus (IFG), superior parietal lobule (SPL), lateral occipital complex (LOC) and finally early visual cortex (EVC). **C)** The suppression points in INS and IFG appeared significantly earlier than the suppression points in EVC and LOC. a.u. = arbitrary units.

In summary, in line with the RT results, the ROI and whole-brain results suggested that neural effects of contextual expectation emerged progressively within a context and reflected activity in sensory and downstream regions involved in visual object prediction (Manahova et al., 2018; Richter et al., 2018; Richter and de Lange, 2019; He et al., 2022).

### Distinct temporal profiles of contextual expectation suppression across the cortical hierarchy

Finally, we characterized when expectation suppression first emerged for each ROI by computing the bin value corresponding to the first zero-crossing of the expectation suppression profile (hereafter termed ‘suppression point’). The suppression point represented when BOLD responses to expected image pairs first became suppressed relative to unexpected ones over the course of a context block. First, the across-participants’ suppression point (s, mean ± SEM) was significantly above zero for all ROIs: EVC (s = 5.64 ± 0.02, t_(33)_ = 8.65, p < 0.001, d_z_ = 1.48), LOC (s = 5.39 ± 0.02, t_(33)_ = 8.70, p < 0.001, d_z_ = 1.49), SPL (s = 4.81 ± 0.05, t_(33)_ = 2.78, p = 0.009, d_z_ = 0.48), IFG (s = 3.50 ± 0.02, t_(33)_ = 4.71, p < 0.001, d_z_ = 0.81) and INS (s = 2.85 ± 0.02, t_(33)_ = 3.99, p < 0.001, d_z_ = 0.68). In line with the ROI and whole-brain results described above, these results are compatible with a consistent carry-over effect of expectation suppression for the expected pairs of the previous block, which subsided as the system attuned itself to the current context. Crucially, as shown in Figure 4B, different suppression points appeared across the cortical hierarchy: first in the insula (INS), followed by the remaining downstream regions (IFG, SPL), and finally in the ventral visual stream (LOC, EVC). In particular, the suppression points in INS and IFG appeared significantly earlier than the suppression points in EVC and LOC (Figure 4C), as indicated by the respective planned paired t tests (Table 4). Importantly, we confirmed that these results are not dependent on the mask size of the ROIs: the direction and statistical significance of all effects were identical across all mask sizes between 100 and 400 voxels. In sum, distinct profiles of contextual expectation suppression emerged across the cortical hierarchy, with surprise signals first being evident in the insula and in frontoparietal regions, and later arising in sensory regions. These results suggest that downstream regions may guide the selection of appropriate sensory expectations in a volatile environment, where different contingencies appear depending on context.

**Table 4.** Results of suppression point analysis in the categorization task. Planned paired t tests comparing the jackknife-estimated suppression points of sensory ROIs (EVC, early visual cortex; LOC, lateral occipital complex) versus downstream ROIs (SPL, superior parietal lobule; IFG, inferior frontal gyrus; INS, insula). Prior to testing for statistical significance, t-values were transformed to *t_corrected_* = t/(n – 1).

| ROI comparison | $t_{corrected}$ (33) | $p$ | $d_z$ |
| --- | --- | --- | --- |
| EVC vs SPL | 0.49 | 0.627 | 0.08 |
| EVC vs IFG | 2.66 | 0.012 | 0.46 |
| EVC vs INS | 3.14 | 0.003 | 0.54 |
| LOC vs SPL | 0.38 | 0.707 | 0.06 |
| LOC vs IFG | 2.60 | 0.014 | 0.45 |
| LOC vs INS | 2.80 | 0.008 | 0.48 |

### No activity modulation for task-irrelevant contextual expectations

A top-down gating mechanism for the selection of contextual expectations may critically depend on task relevance. In the categorization task, predictions were in fact task-relevant: knowing which trailing images most likely followed each leading image served the categorization process. We addressed whether we found the same neural effects of contextual expectation in a separate oddball task where expectations were instead task-irrelevant: oddballs were fully counterbalanced with respect to image status (leading / trailing) and category (electronic / non-electronic), thus object predictions did not serve oddball detection.

We anticipated participants to be highly correct at detecting the oddballs, since we pre-selected them based on accurate performance (see section ‘Exclusion criteria - fMRI’). Indeed, the across-participants’ proportion of correct responses was close to ceiling (0.96 ± 0.01, mean ± SEM), which confirmed high vigilance during the oddball task. Crucially, here we did not find any expectation suppression throughout the brain, in sharp contrast with the results of the categorization task (Figure 5A). In particular, while there was no significant interaction between expectation and context half in any ROI, we observed a significant main effect of expectation in LOC and SPL, with qualitatively similar profiles across ROIs (Figure 5B and Table 5). Furthermore, the whole-brain analysis did not show any significant brain activations for the main effect of expectation, the main effect of context half and the interaction between them (Figure 5C).

**Figure 5.**
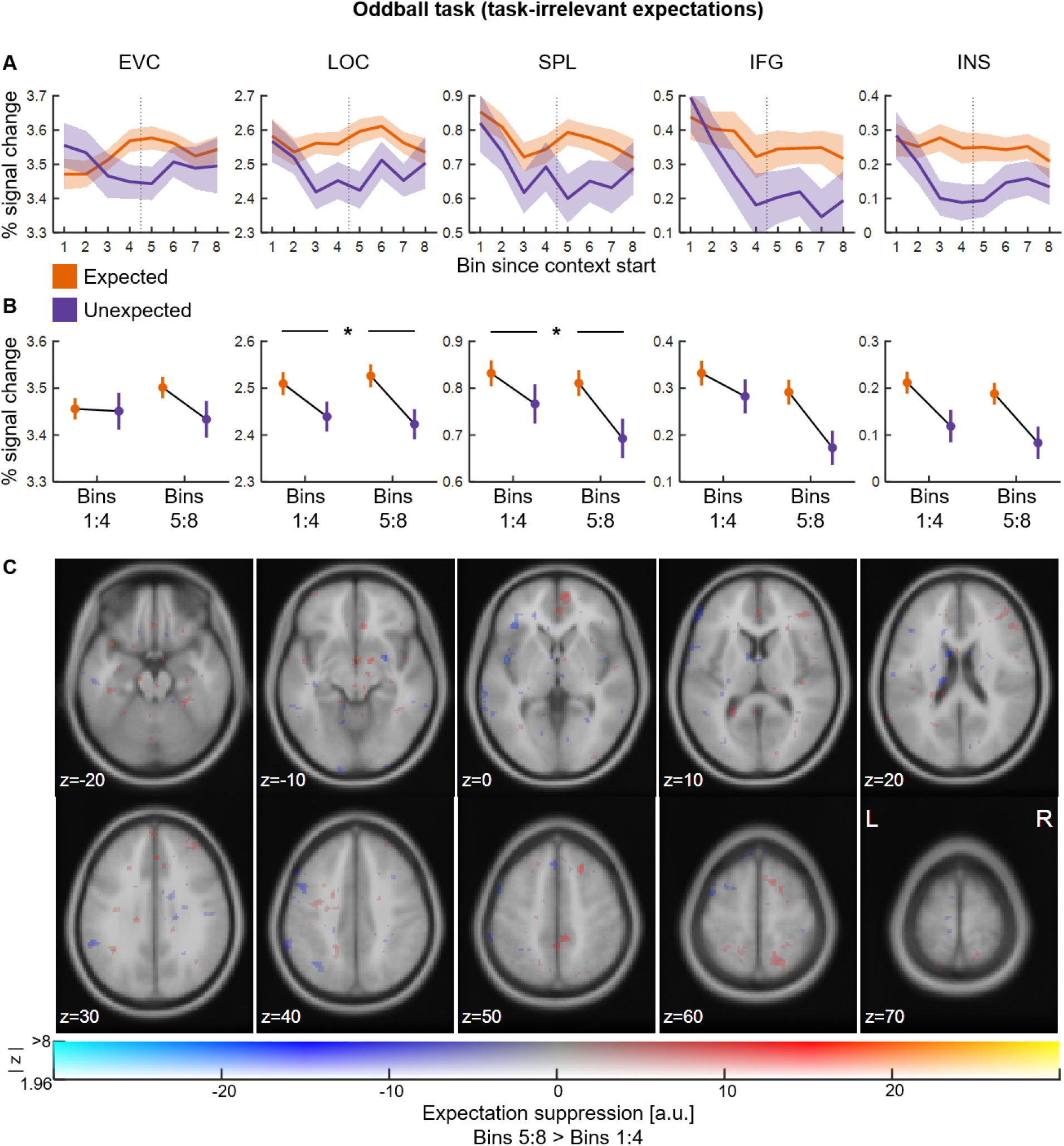
Neural effects of contextual expectation in the oddball task. **A)** Across participants’ mean (± SEM) % signal change relative to baseline for each ROI as function of expectation (expected: orange; unexpected: purple) and bin (1-8) since the start of a context block. **B)** Same data plotted as a function of expectation and context half (first half, bins 1:4 / second half, bins 5:8). **C)** Expectation suppression (i.e. unexpected > expected) in the second context half (bins 5:8) relative to the first context half (bins 1:4), overlaid on the MNI152 2mm template. Colours represent the parameter estimates: red-yellow clusters reflect expectation suppression; opacity represents the associated z statistics. We did not observe expectation suppression neither in the ROI analysis nor at the whole-brain level. a.u. = arbitrary units, * p < 0.05.

**Table 5.** Results of expectation × context half ROI analysis in the oddball task. Main effects and interactions of the 2 (expectation: expected / unexpected) × 2 (context half: bins 1:4 / bins 5:8) repeated measures ANOVA on % signal change relative to baseline for each ROI: EVC, early visual cortex; LOC, lateral occipital complex; SPL, superior parietal lobule; IFG, inferior frontal gyrus; INS, insula.

| ROI | $F_{(1,33)}$ | $p$ | $\eta_p^2$ |
| --- | --- | --- | --- |
| **EVC** |  |  |  |
| expectation | 0.56 | 0.458 | 0.02 |
| context half | 0.08 | 0.781 | 0.01 |
| expectation × context half | 0.63 | 0.432 | 0.02 |
| **LOC** |  |  |  |
| expectation | 4.17 | 0.049 | 0.11 |
| context half | 0.00 | 0.996 | 0.00 |
| expectation × context half | 0.20 | 0.661 | 0.01 |
| **SPL** |  |  |  |
| expectation | 5.59 | 0.024 | 0.14 |
| context half | 0.62 | 0.438 | 0.02 |
| expectation × context half | 0.49 | 0.487 | 0.01 |
| **IFG** |  |  |  |
| expectation | 3.41 | 0.074 | 0.09 |
| context half | 2.04 | 0.163 | 0.06 |
| expectation × context half | 0.93 | 0.342 | 0.03 |
| **INS** |  |  |  |
| expectation | 3.98 | 0.054 | 0.11 |
| context half | 0.38 | 0.541 | 0.01 |
| expectation × context half | 0.03 | 0.866 | 0.00 |

Complementarily, we tested for the emergence of expectation suppression over the course of the 8 individual context bins in all ROIs. After fitting a linear regression to the expectation suppression profile per participant and ROI, the across-participants’ estimated regression slope (b, mean ± SEM) was not significantly different from zero for any ROIs, with all Bayesian results confirming moderate evidence for the absence of an effect: EVC ( b = −1.03 ± 1.93, t_(33)_ = −0.53, p = 0.596, d_z_ = −0.09, B_01_ = 4.76), LOC ( b = 0.09 ± 1.68, t_(33)_ = 0.05, p = 0.959, d_z_ = 0.01, B_01_ = 5.56), SPL (b = −0.04 ± 1.59, t_(33)_ = −0.02, p = 0.982, d_z_ = −0.01, B_01_ = 5.56), IFG ( b = −1.81 ± 1.99, t_(33)_ = −0.91, p = 0.368, d_z_ = −0.16, B_01_ = 3.70) and INS (b = −0.82 ± 1.47, t_(33)_ = −0.56, p = 0.579, d_z_ = −0.10, B_01_ = 4.76). Once again, the direction and statistical significance of all effects were identical across all mask sizes between 100 and 400 voxels.

Collectively, the ROI and whole-brain results showed evidence against expectation suppression throughout the brain in the case of task-irrelevant contextual expectations, suggesting that task relevance may represent a boundary condition for the top-down gating of contextual expectations.

## Discussion

The present study examined the behavioural and neural effects of contextual sensory expectation as observers updated their beliefs in a volatile environment. After developing visual object predictions conditioned on context, participants showed progressively faster responses to expected than unexpected stimuli within the context, suggesting that they used their predictive knowledge to optimise task performance. This was accompanied by progressively suppressed BOLD responses for unexpected compared to expected trials (i.e. expectation suppression) across three widespread systems: in early visual cortex, lateral occipital complex and fusiform gyrus, which are part of the ventral visual stream for object processing; in superior parietal lobule, intraparietal sulcus and inferior frontal gyrus, which are central nodes of the frontoparietal network for top-down attention gating; in the insula and anterior cingulate cortex, which belong to the salience network, a control hub for surprise detection and cognitive control. Importantly, we found distinct neural profiles across the cortical hierarchy: expectation suppression appeared first in the insula and frontoparietal regions, followed by the ventral visual stream and early visual cortex. Finally, this neural pattern pertained selectively to task-relevant predictions. As a whole, the present study suggests that an insular and frontoparietal executive control network may gate access to task-relevant contextual expectations to guide behaviour in a complex and dynamic environment, where changing contexts determine the sensory contingencies we experience.

### Encoding and retrieval of contextual sensory expectations

To investigate how sensory expectations are flexibly deployed conditioned on context, we created a volatile environment with two state spaces hierarchically controlled by an independent contextualizing signal, consistent with structure learning paradigms (Collins and Frank, 2013; Gershman, 2017; Collins, 2018). Across the main fMRI experiment and a supplementary structure learning experiment, participants successfully encoded and retrieved contextual regularities after they were instructed to discover structure conditioned on context. These findings support and extend previous work on structure learning (Collins and Frank, 2013; Gershman, 2017; Collins, 2018) by showing that, also outside the realm of reinforcement learning, organizing knowledge by context and focusing selectively on the task-relevant sensory contingencies scaffolds fast and robust learning inside a changing environment (Wilson and Niv, 2012). In striking contrast, in an incidental exposure experiment matched in terms of duration and environmental structure, participants did not show any evidence of learning neither in terms of implicit RT facilitation nor in terms of explicit recognition. Thus, the incidental and automatic extraction of regularities, known as statistical learning (Saffran et al., 1996; Turk-Browne et al., 2010; Batterink and Paller, 2019; Batterink et al., 2019), appears to be at least slowed down and hence not manifest in a hierarchically structured and volatile environment like the one employed in the present study (see also de Waard et al., 2022 for complementary evidence during statistical learning of distractor suppression). Such evidence has theoretical implications for statistical learning, which is typically described as a rapid mechanism able to extract environmental regularities in a matter of few exposures (Aslin, 2017; Saffran and Kirkham, 2018; Sherman et al., 2020). Traditional statistical learning experiments present participants with fixed environments that contain simple regularities, such as chunks of stimuli linked by deterministic dependencies (Saffran et al., 1996; Turk-Browne et al., 2009; Henin et al., 2021), which arguably represent an oversimplification of real-life conditions. Increasing structure complexity, here in the form of context-dependent, probabilistic and volatile regularities, clearly hinders the effectiveness of statistical learning and hence highlights a possible boundary condition that deserves further investigation, as recently discussed (Conway, 2020).

### Updating sensory expectations in a volatile environment

Having established robust structure learning in the fMRI experiment, we investigated its behavioural and neural consequences using response times and voxel-wise BOLD responses. Both measures showed that contextual expectation effects emerged progressively in a categorization task where predictions were task-relevant. This pattern of results may look surprising, given that a preliminary explicit cue was fully informative of the upcoming context identity and its associated sensory contingencies. Crucially, our volatile environment required the inhibition of irrelevant contextual contingencies when the context changed, hence generating switching costs at the beginning of a new context period (Monsell, 2003; Koch et al., 2010; Collins, 2017). This was substantiated by a consistent carry-over effect of expectation suppression for the expected pairs of the previous block, which subsided as the system attuned itself to the current context.

Brain areas showing progressive contextual expectation suppression within a context comprised sensory and downstream regions involved in visual object prediction (Richter et al., 2018; Richter and de Lange, 2019; He et al., 2022). Crucially, distinct neural profiles appeared across the cortical hierarchy. An insular and frontoparietal system first reacted to expectation violations. On the one hand, the insula is a central node of the salience network, a control hub that marks relevant events in conflict with current states (Menon and Uddin, 2010; Uddin, 2015). In particular, it reflects surprise both during value-based decision-making and during sensory processing (Fazeli and Büchel, 2018; Loued-Khenissi et al., 2020). On the other hand, the posterior parietal cortex (SPL / IPS in particular) and the inferior frontal gyrus are part of the frontoparietal network for attention (re)orienting (Corbetta and Shulman, 2002; Shomstein and Behrmann, 2006; Corbetta et al., 2008; Scolari et al., 2015). Further, the inferior frontal gyrus is thought to be involved in switching among ‘task sets’ by inhibiting irrelevant task representations when the context changes (Aron et al., 2004; Sakai, 2008; Hyafil et al., 2009). Together, these two systems are known to enable adaptive behaviour by marking salient events and directing processing resources toward them (Shulman et al., 2009; Menon and Uddin, 2010). As previously suggested (Meyer and Olson, 2011; den Ouden et al., 2012; Richter et al., 2018), a stronger neural response to unexpected than expected stimuli fits this computational goal, because unexpected events may instruct the need to update expectations in a volatile environment.

### Origin of contextual sensory expectations in the brain

The presence of distinct neural profiles across the cortical hierarchy, with expectation suppression first evident in the insula and frontoparietal regions, followed by the ventral visual stream and early visual cortex, offers insight into how expectation effects may arise in the brain. The neural mechanisms implementing the effect of expectation on perception likely depend on the specific type of environmental regularities (de Lange et al., 2018). On the one hand, expectations deriving from stable features of the physical environment can become encoded in the tuning properties of our sensory cortices, as a result of long-term phylogenetic plasticity (Teufel and Fletcher, 2020). One example is the overrepresentation of early visual neurons that are tuned to cardinal than oblique orientations, as well as narrower tuning curves for cardinal orientations, which are more frequent in the natural world (Girshick et al., 2011). Further, expectations deriving from long-term semantic memory support visual recognition under high perceptual uncertainty (i.e. low signal-to-noise ratio) via top-down signals from medial prefrontal to early visual cortex (Summerfield et al., 2006; Summerfield and Egner, 2009). On the other hand, contextual expectations arising from arbitrary and volatile contingencies may either emerge from local short-term plasticity (Reber, 2013; Conway, 2020) or require the involvement of downstream executive control regions that monitor the environment for surprising events (Shulman et al., 2009; Menon and Uddin, 2010; Uddin, 2015; Conway, 2020). However, at present, it is unclear whether contextual expectation effects in sensory areas arise from local synaptic plasticity or instead emerge via top-down signals from downstream areas. Previous work is compatible with both hypotheses, as it employed extensive behavioural training before neuroimaging that could have generated synaptic plasticity in sensory areas (Meyer and Olson, 2011; Kaposvari et al., 2018; Richter et al., 2018; Richter and de Lange, 2019; He et al., 2022). On the contrary, the present study employed a very short training period right before scanning. This experimental feature, coupled with the empirical observation that expectation suppression first appeared in downstream regions, suggests that contextual expectation effects may arise in downstream executive control regions and be subsequently fed back to functionally relevant sensory regions (here, the object-selective ventral visual stream). In sum, we suggest that an insular and frontoparietal executive control system, sensitive to surprising events in conflict with current task-relevant predictions (Shulman et al., 2009; Menon and Uddin, 2010; Uddin, 2015; Conway, 2020), may guide the selection of appropriate contextual sensory expectations in a volatile environment, where arbitrary contingencies are hierarchically structured by context. Accordingly, task relevance critically mediated our findings: we observed distinct neural profiles of expectation suppression across the cortical hierarchy only in the case of task-relevant predictions, which could be used to optimise task execution; no such evidence was found in a separate task where predictions were task-irrelevant. These results place important constraints on neurocomputational theories that cast perception as an automatic integration of prior knowledge and sensory information (Rao and Ballard, 1999; Friston, 2005, 2010). Interestingly, the present results also extend work showing that a frontoparietal network encompassing inferior frontal gyrus and posterior parietal cortex is responsible for the selection of task-relevant sensory expectations in a multidimensional environment, where different contingencies co-exist (Niv et al., 2015; Leong et al., 2017). Collectively, the top-down gating of task-relevant sensory expectations from downstream executive control areas to sensory areas may enable adaptive behaviour in our complex and dynamic world.

## Acknowledgements

The present work was supported by the EC Horizon 2020 Program (ERC Consolidator Grant “Surprise” 101000942 to F.P.d.L). The authors thank Yamil Vidal and Jakub Szewczyk for helpful comments on a previous version of the manuscript.

